# Contractile acto-myosin network on nuclear envelope remnants positions human chromosomes for mitosis

**DOI:** 10.1101/459750

**Authors:** Alexander JR Booth, Zuojun Yue, John K Eykelenboom, Tom Stiff, GW Gant Luxton, Helfrid Hochegger, Tomoyuki U Tanaka

## Abstract

To ensure proper segregation during mitosis, chromosomes must be efficiently captured by spindle microtubules and subsequently aligned on the mitotic spindle. The efficacy of chromosome capture by the mitotic spindle can be influenced by how widely chromosomes are scattered in space. However it is unknown whether chromosome scattering is regulated, prior to the interactions with spindle microtubules, in early mitosis. Here, we quantify chromosome-scattering volume (CSV) and find that it is reduced immediately after nuclear envelope breakdown (NEBD) in human cells. The reduction of CSV occurs independently of microtubules and is therefore not an outcome of interactions between chromosomes and the spindle. We find that, prior to NEBD, an acto-myosin network is assembled in a LINC complex-dependent manner on the cytoplasmic surface of the nuclear envelope. This acto-myosin network remains on the nuclear envelope remnant soon after NEBD, and its myosin-II-mediated contraction reduces CSV and facilitates chromosome interactions with spindle microtubules. Thus we find a novel mechanism that positions chromosomes in early mitosis to ensure efficient chromosome–spindle interactions.

## Introduction

The mitotic spindle is composed of microtubules (MTs) and plays central roles in chromosome segregation during mitosis. To ensure proper chromosome segregation, chromosomes must be efficiently captured and aligned on the mitotic spindle prior to their segregation (*1, 2, 3*). For this, chromosomes should establish initial interaction with spindle MTs soon after nuclear envelope breakdown (NEBD). Subsequently, sister chromatids must interact with MTs from opposite spindle poles (chromosome bi-orientation). Despite being an active area of investigation, the cellular and molecular mechanisms promoting chromosome interactions with the mitotic spindle remain incompletely defined.

The efficiency of establishing chromosome interaction with spindle MTs is likely to be significantly affected by the positioning of chromosomes relative to the spindle poles immediately following NEBD. For example, if chromosomes were located far away from the spindle poles, more time would be required to establish the initial chromosome interaction with spindle MTs. Moreover, should chromosomes be located behind a spindle pole, i.e. opposite from the bulk of spindle and outside of the pole-to-pole region, their interaction with MTs from the other spindle pole would be delayed.

### Chromosome scattering volume is reduced in early prometaphase, independently of spindle MTs

To investigate chromosome positioning during early mitosis, we quantified how widely chromosomes were scattered in space. To do this, we acquired three-dimensional images of fluorescently labelled DNA (SiR-DNA) in live mitotic human U2OS cells. Subsequently, we obtained the convex hull of the chromosome distribution. The covex hull in two and three dimension (2D and 3D) is the minimal polygon and polyhedron, respectively, that wrap multiple objects (chromosomes in this case). This is schematically shown in Figure 1A, using a shrinking ‘rubberband’ and ‘balloon’ for intuitive demonstration. We determined the volume of this minimal polyhedron or shrunk balloon that wrapped chromosomes, and defined it as the chromosome scattering volume (CSV), which represents how widely chromosomes scatter in 3D space.

**Figure 1.**
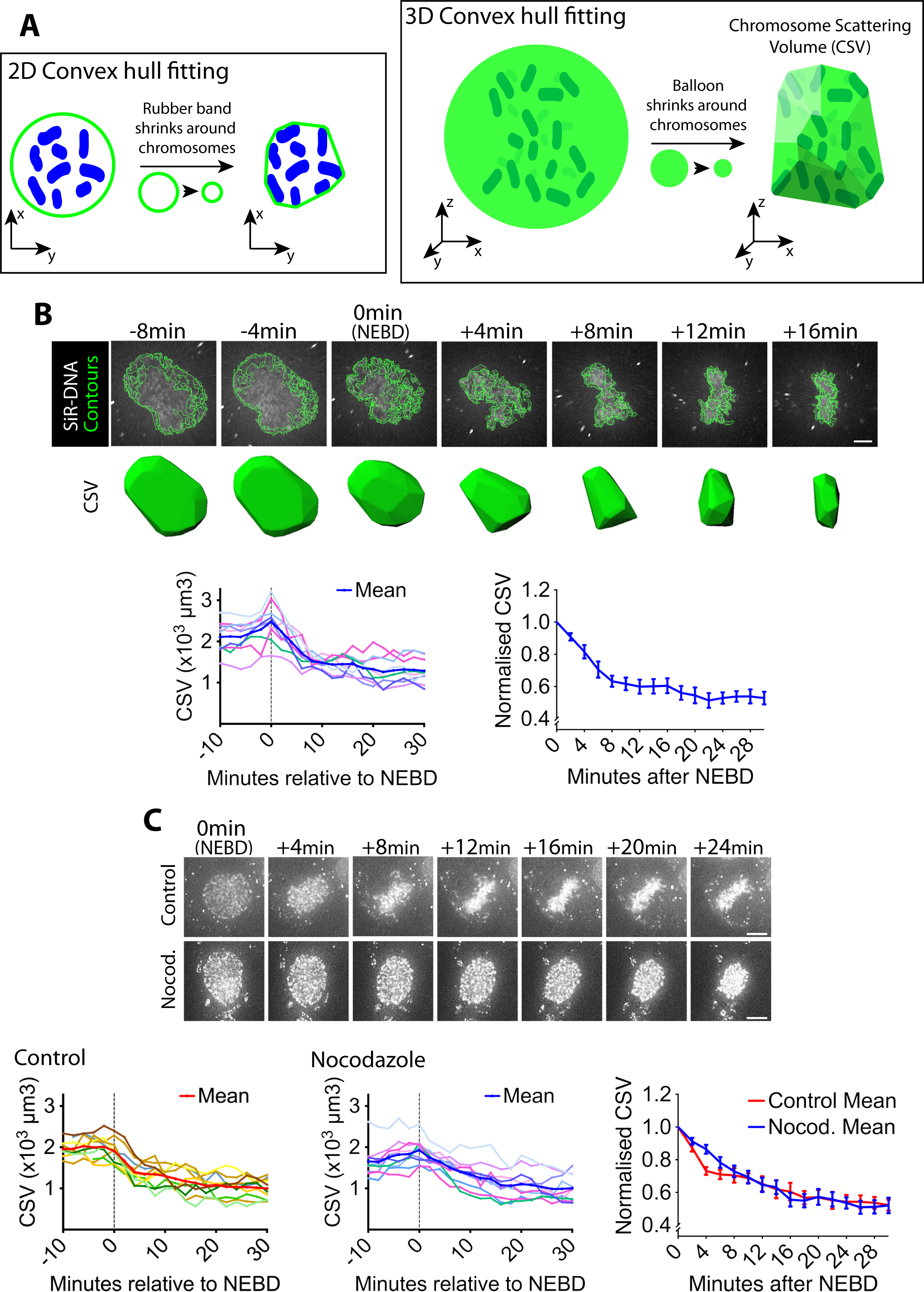
Chromosome scattering volume is reduced in early prometaphase, independently of spindle MTs. A) Diagrams show how the chromosome scattering volume (CSV) was defined. Left-hand diagram shows how the convex hull was generated in two dimensions (2D) to represent chromosome distribution. A ‘rubber band’ (green) shrinks around chromosomes (blue) in 2D to create a convex hull or a minimum polygon wrapping chromosomes. Right-hand diagram shows how the convex-hull was generated in three dimensions (3D) to represent chromosome distribution. A ‘balloon’ (green) shrinks around chromosomes (dark green) in 3D to create a convex hull or a minimal polyhedron wrapping chromosomes. The volume of the 3D convex hull (chromosome scattering volume; CSV) quantifies how widely chromosomes are scattered in space. B) CSV decreases immediately after NEBD. Images (top) are z-projections (z-sections are projected to 2D images) of a representative cell stained with SiR-DNA to visualise chromosomes (white) alongside their perimeter contours at z-sections (green lines; see Materials and Methods). Time is shown, relative to NEBD. Timing of NEBD was determined as in Figure S1A. Scale bars, 6µm. Bottom shows corresponding CSV (green). The left-hand graph shows CSV measurements in individual cells aligned by the time relative to NEBD (n=9). To make the right-hand graph, these CSV values were normalized to the CSV value at NEBD in each cell, and the mean of the normalized CSV values were plotted at each time point. Error bars, s.e.m. C) CSV decreases after NEBD even in the absence of MTs. Images (z-projections) show representative cell with SiR-DNA-stained chromosomes, which entered mitosis in the presence of 3.3µM nocodazole (Nocod., bottom) or DMSO (control, top). 0.5 µM MK-1775 Wee1 inhibitor was used in both conditions to allow cells to enter mitosis. Time is shown relative to NEBD, whose timing was determined as in Figure S1A. Scale bars, 10µm. Left-hand and center graphs show CSV measurements in individual control and nocodazole-treated cells, respectively (n=10 each). The right-hand graph compares the means of normalized CSV between the nocodazole-treated and control cells (error bars, s.e.m.), as in the right-hand graph in B. Reduction of normalized CSV was not significantly different between control and nocodazole treatment, when all the time points were considered by two-way ANOVA (*p*=0.81). Nonetheless, normalized CSV was significantly different between the two groups at +4 min (t-test, *p*=0.0016), but not at other time points.

To synchronize U2OS cells in early mitosis, we used a previously-described U2OS cell line in which the *CDK1* gene was replaced, by genome engineering, with *cdk1-as*, whose gene product Cdk1-as can be specifically inhibited by an ATP analogue 1NM-PP1 (*4*). The U2OS *cdk1-as* cells were arrested in G2 with 1NM-PP1, and subsequently released by 1NM-PP1 washout to synchronously undergo mitosis. NEBD was identified by the release of the nuclear-localizing fluorescent reporter (GFP-LacI-NLS) into the cytoplasm (Figure S1A). Using these methods, we quantified the CSV in synchronized U2OS *cdk1-as* cells as they progressed from NEBD (defined as t = 0) into prometaphase. We found that CSV was prominently reduced within the first 8 min following NEBD, and reduction continued more slowly over the following ∼10 min (Figure 1B). CSV reduction was also observed in asynchronous, wild-type U2OS cells (Figure S1B), indicating that it was not an artefact caused by *cdk1-as*-mediated cell cycle synchronization. Consequently, we used U2OS *cdk1-as* cells for synchronous entry into mitosis for the rest of the experiments presented in this work, unless otherwise stated.

It is possible that the CSV reduction observed following NEBD was an outcome of the interaction between chromosomes and spindle MTs. To test this possibility, we quantified CSV in cells treated with nocodazole. After nocodazole treatment, MTs were almost completely depleted (Figure S1C). In control cells, chromosomes moved inward after NEBD, and subsequently aligned on the metaphase plate (Figure 1C, top-row images). In cells lacking visible MTs, chromosomes also moved inward after NEBD, but then remained in a spherical formation (Figure 1C, bottom-row images). The CSV was reduced with very similar kinetics in both the presence and absence of MTs (Figure 1C, graphs). CSV was slightly smaller at 4 min following NEBD in control cells; this could be due to mild compression of the nuclear envelope (NE) remnants caused by the rapid MT-dependent inward movement of centrosomes (Figure 1C, image at +4 min in control). Consequently, we conclude that the overall CSV reduction observed following NEBD is not an outcome of chromosome interaction with spindle MTs.

Alternatively, chromosome condensation, which occurs in early mitosis, may cause the CSV reduction after NEBD. To test this possibility, we quantified the total volume of chromosome mass (chromosome mass volume [CMV]; Figure S1D) after NEBD. Unlike the CSV, the CMV did not considerably change following NEBD. Thus, we conclude that the reduction in CSV was not due to chromosome condensation, which likely occurs mainly before NEBD (*5*).

### Actin accumulates outside of the NE in prophase in a LINC complex-dependent manner, and the actin network shrinks after NEBD

What mechanisms, then, might promote CSV reduction after NEBD? The CSV reduction could be explained if chromosomes were surrounded by a physical barrier that contracts following NEBD. Consistent with this model, we found that in early mitosis an actin network accumulated on the NE, peaking in intensity around NEBD and persisting after it (Figure 2A). The volume inside this actin network was rapidly reduced within 6 min following NEBD (Figure S2A), suggesting the contraction of the network. During this time, the chromosomes were surrounded by, and the outermost chromosomes were found immediately adjacent to, the actin network (Figure S2B). We used *cdk1-as* in these experiments to synchronize cells, however the actin network was also observed in wild-type U2OS cells (Figure S1B, yellow arrowheads), indicating that it is not an artefact due to *cdk1-as* regulation.

**Figure 2.**
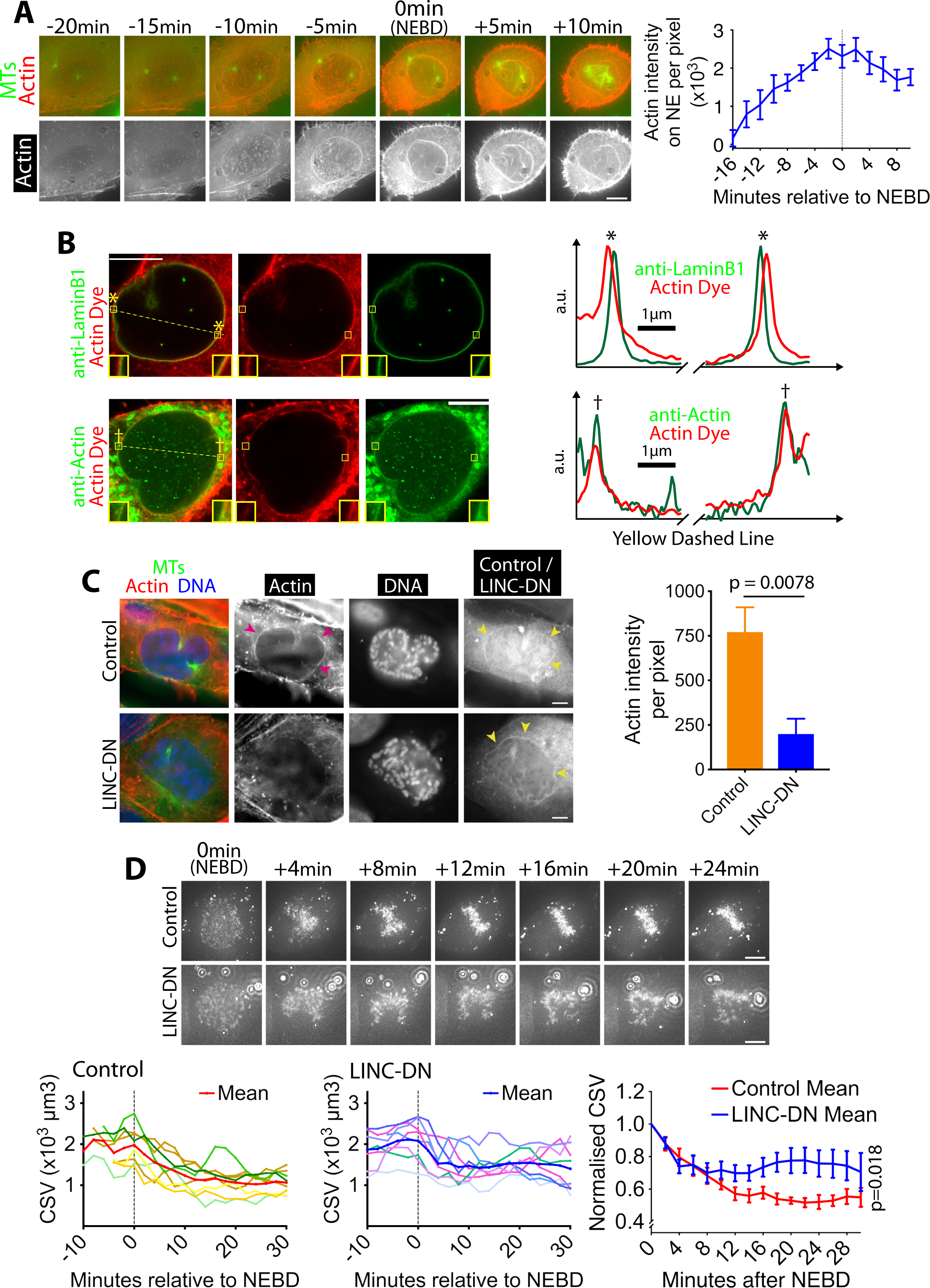
Actin accumulates outside of the NE in prophase, and its network shrinks after NEBD. A) Actin accumulates on the NE around NEBD. Images (z-projections) show a representative cell expressing GFP-tubulin and mCherry-Lifeact (that fluorescently marks F-actin). Time is relative to NEBD. Timing of NEBD was determined by the influx of cytoplasmic GFP-tubulin into the nucleus. Scale bars, 10µm. Graph shows mean Lifeact fluorescence intensity (per pixel) around the nucleus over time (n=8; error bars, s.e.m). B) Actin localizes outside of the NE. Images show single super-resolution z-sections of cells, which were fixed and stained for actin with phalloidin (actin dye; red). Cells were also immunostained (green) with anti-Lamin B1 (top) or anti-actin antibody (bottom), respectively. The same secondary antibody was used for the two immuno-stainings. These cells were undergoing chromosome compaction (confirmed by DNA staining). Insets show magnification of the regions in yellow boxes. Scale bar, 10µm. Graphs show intensity of actin dye (red) and immunostaining (green) along the dashed yellow lines in left images (middle parts are omitted). The peaks in graphs, marked with asterisks (top) and daggers (bottom), locate at the regions in yellow boxes in left images. a.u, arbitrary unit. C) The LINC complex is required for accumulation of the actin network. Images show single z-sections of representative cells, expressing either an RFP-tagged LINC-DN construct (SR-KASH) or an RFP-tagged control construct (KASHΔL) (*9*). Cells were stained for actin with phalloidin (red), DNA with Hoechst 33342 (blue) and MTs with anti-tubulin antibody (green). Right-hand-most images show RFP signals showing localisation of the LINC-DN or control construct. To focus on cells in prophase or early prometaphase, we analyzed cells where chromosome compaction had started and the centrosomes had separated, but bipolar spindle formation had not yet been completed. 0.5 µM MK-1775 Wee1 inhibitor was used to allow cells to enter mitosis in both conditions. Magenta arrowheads indicate the actin network on NE in control. Yellow arrowheads indicate localization of LINC-DN and its control on NE. Scale bars, 10µm. Graph on right shows mean intensity of actin signal around the nucleus in cells expressing LINC-DN vs a control construct (control n=18, LINC-DN n=10). *p* value was obtained by *t*-test. Error bars, s.e.m. D) Removal of the actin network results in slower CSV reduction after NEBD. Images (z-projections) show a cell with SiR-DNA-stained chromosomes, expressing either LINC-DN or a control construct. Localization of LINC-DN and its control was as C. MK-1775 was used in both conditions as in C. Time is relative to NEBD, whose timing was determined as in Figure S1A. Scale bars, 10µm, Left-hand and center graphs show CSV measurements in individual cells expressing control and LINC-DN constructs, respectively (control n=7, LINC-DN n=8). The right-hand graph compares the means of normalised CSV (as in Figure 1B) Error bars, s.e.m. *p* value (control vs LINC-DN) was obtained by two-way ANOVA.

We investigated in more detail the localization of the actin network on the NE or its remnants, following NEBD. Using the Airyscan super-resolution microscopy, we compared the localization of the actin network and lamin B1; the latter is found on the nucleoplasmic surface of the NE (*6*) and remains there soon after NEBD (*7*). The actin network was separated from lamin B1 by 150–200 nm (Figure 2B), suggesting that it may localize on the cytoplasmic surface of the NE or its remnants following NEBD. A candidate receptor for the actin network on the NE is the LINC (linker of nucleoskeleton and cytoskeleton) complex, one end of which is anchored on the NE while the other end extends to the cytoplasm and interacts with actin (*8*). To test whether the NE-association of the actin network during early mitosis was LINC complex-dependent, we visualized the actin network in cells expressing a previously-described dominant negative construct of the LINC complex (LINC-DN) or its control (SR-KASH and KASHΔL, respectively (*9*)). As expected, both constructs localized to the NE. (Figure 2C, yellow arrowheads). Crucially, the fluorescence intensity of the actin network was significantly diminished in cells expressing LINC-DN, compared with cells expressing its control (Figure 2C, magenta arrowheads, graph). Therefore the actin network localizes to the cytoplasmic surface of the NE and its remnants, following NEBD, in a LINC complex-dependent manner.

If the actin network were a physical barrier, confining chromosomes, and its contraction were to lead to the CSV reduction after NEBD, LINC complex disruption should also alleviate CSV reduction. We tested this hypothesis by measuring the CSV in cells expressing LINC-DN and its control (Figure 2D) and found that the CSV reduction indeed slowed after NEBD with the LINC-DN expression, compared with the control expression. Thus, it is likely that the actin network on NE remnants confines chromosomes and that its contraction leads to CSV reduction.

### Myosin II activity is required for actin network contraction and CSV reduction after NEBD

We next addressed how the actin network contracts on the NE remnants to reduce the CSV after NEBD. We first tested whether actin depolymerization causes the CSV reduction. An actin depolymerization inhibitor jasplakinolide had no effect on CSV reduction after NEBD (Figure S3A), though it successfully inhibited subsequent cytokinesis as previously reported (*10*). Thus it is unlikely that actin depolymerization promotes actin network contraction or CSV reduction.

The actin-dependent motor protein myosin II is a major mediator of acto-myosin contractility in non-muscle cells (*11*). Thus, we next tested the possible involvement of myosin II in the actin network contraction and CSV reduction. We found that myosin II co-localized with the actin network on the NE (Figure S3B, yellow arrowheads). We next inhibited the myosin II activity using para-nitroblebbistatin (pnBB), a photo-stable version of its inhibitor blebbistatin, which is suitable for live-cell imaging (*12*). Notably, pnBB treatment alleviated both the CSV reduction (Figure 3A) and the actin network contraction (Figure S3C) following NEBD. While pnBB did not significantly alter the distance between spindle poles soon after NEBD (Figure S3C), it often caused the actin network to extend beyond the spindle poles at the initiation of spindle formation (Figure S3D, pnBB, 5–20 min). The pnBB treatment did not affect CMV (see Figure S1D) after NEBD (Figure S3E), suggesting that while myosin II activity does not change chromosome condensation, it does facilitate CSV reduction. Moreover, the pnBB treatment alleviated the CSV reduction in the presence of nocodazole (Figure 3B), indicating that myosin II promotes the actin network contraction and CSV reduction after NEBD in a spindle MT-independent manner. Double treatment with pnBB and Nocodazole seemed to further lessen the CSV reduction during 0–6 min, compared with pnBB treatment alone (compare Figure 3A and 3B). Thus a MT-dependent mechanism, leading to mild NE remnant compression by centrosomes (see Figure 1C), and a myosin II-dependent mechanism may work in concert to reduce CSV during this period.

**Figure 3.**
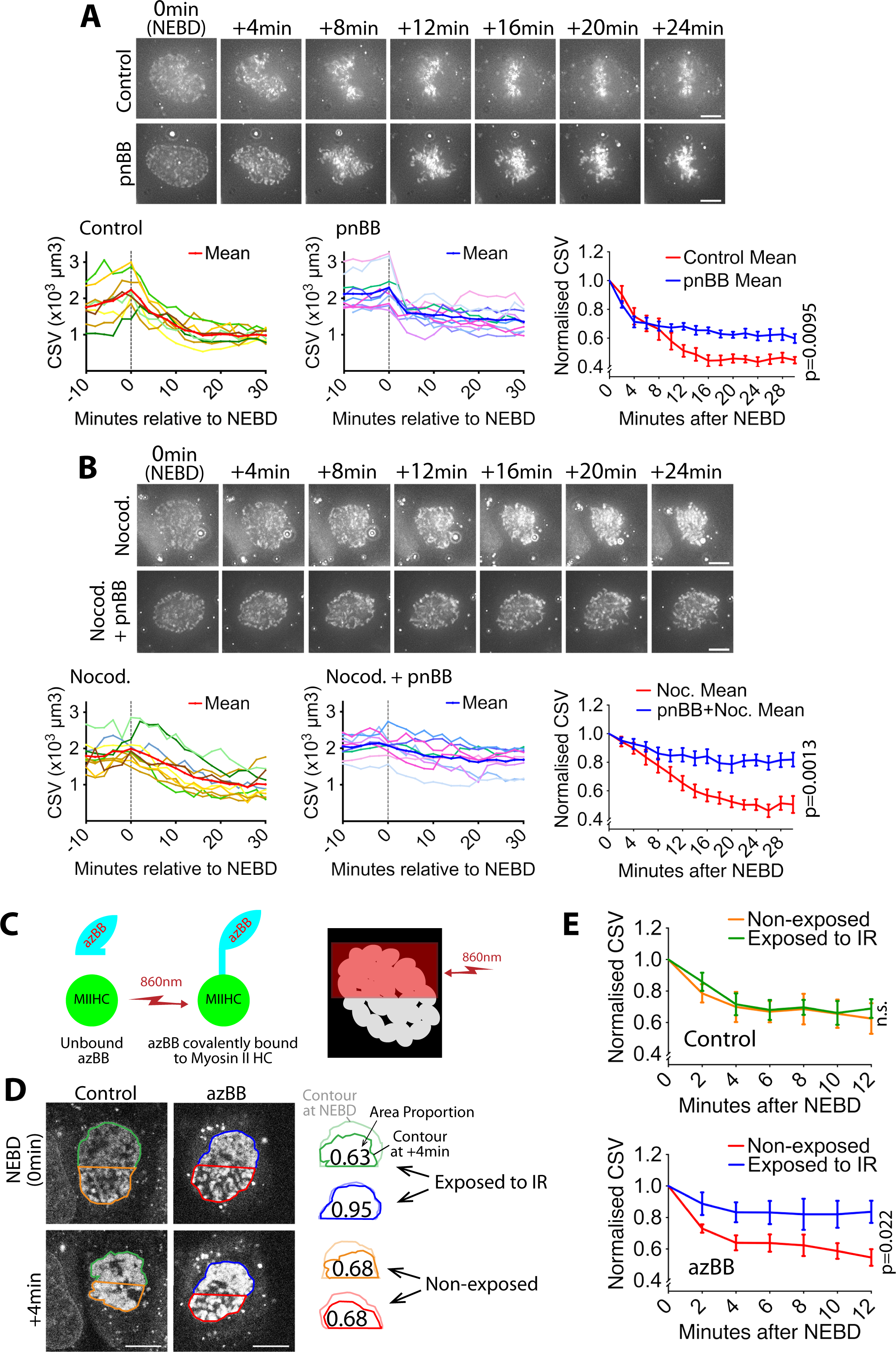
Myosin II activity is required for the CSV reduction after NEBD. A) Inhibition of myosin II activity alleviates CSV reduction after NEBD. Images (z-projections) show representative cells with SiR-DNA-stained chromosomes, which entered mitosis in the presence of 50µM paranitroblebbistatin (pnBB, bottom) or DMSO (control, top). Time was relative to NEBD, whose timing was determined as in Figure S1A. Scale bars, 10µm. Left-hand and center graphs show CSV measurements in individual control and pnBB-treated cells, respectively (pnBB, n=10; control, n=8). The right-hand graph compares the means of normalized CSV (as in Figure 1B) in each condition. Error bars = s.e.m. *p* value (control vs pnBB) was obtained by two-way ANOVA. B) CSV reduction is almost completely alleviated when myosin II activity is inhibited in the absence of MTs. Images (z-projections) show representative cells with SiR-DNA-stained chromosomes, which entered mitosis in the presence of 3.3µM nocodazole plus 50µm pnBB (Nocod. + pnBB, bottom), or 3.3µM nocodazole (Nocod., top). 0.5 µM MK-1775 Wee1 inhibitor was used to allow cells to enter mitosis in both conditions. Time was relative to NEBD, whose timing was determined as in Figure S1A. Scale bars, 10µm. Left-hand and center graphs show CSV measurements in individual cells treated by nocodazole alone and by nocodazole plus pnBB, respectively (nocodazole treated n=10, nocodazole and pnBB treated n=9). The right-hand graph compares the means of normalized CSV (as in Figure 1B) in each condition. Error bars = s.e.m. *p* value (nocodazole vs pnBB+Nocodazole) was obtained by two-way ANOVA. C, D, E) Local inhibition of myosin II activity results in local alleviation of CSV reduction. C) Diagram shows that azidoblebbistatin (azBB) is covalently linked to the myosin II heavy chain and inhibits its activity in the half of the nucleus exposed to infrared light. D) Images (projections of three z-sections) show SiR-DNA-stained chromosomes in representative cells that were incubated in the presence of 5µM azidoblebbistatin (azBB, right) or DMSO (control, left). The half of the nucleus in blue and green was exposed to 860 nm light just prior to NEBD, while the half in red and orange was not. Time was relative to NEBD, whose timing was determined as in Figure S1A. Scale bars, 10µm. Diagrams show how the indicated area was reduced at + 4 min, relative to the area at NEBD. E) Graphs show means of normalized CSV in half of the nucleus, which was exposed vs non-exposed to infra-red (IR), in the presence (bottom) and absence (top) of azBB (control n=5, azBB n=7). Colors of lines match the colors that border the half nuclei in D. *p* value (exposed vs non-exposed to IR) was obtained by two-way ANOVA. n.s., not significant. Error bars, s.e.m. CSV was normalized as in Figure 1B.

A straightforward interpretation of these results is that myosin II within the actin network causes its contraction and the CSV reduction after NEBD. To obtain more evidence for this model, we performed light-induced local inhibition of myosin II activity in mitotic cells. We used azido-blebbistatin (azBB), which covalently binds the heavy chain of myosin II and inhibits its ATPase activity when exposed to infra-red light (*13*). A low concentration of azBB was used to ensure that myosin II was inhibited only at the region exposed to infra-red light. We exposed a half of the nucleus to infra-red light (Figure 3C) and compared the CSV over time in the exposed and non-exposed halves of the nucleus. In the exposed half of the nucleus (Figure 3D, E; blue), the CSV reduction was alleviated, compared to the non-exposed half (red). In the cells not treated with azBB (control), there was no significant difference in CSV between exposed (Figure 3D, E; green) and non-exposed (orange) halves of nuclei. These results support a model in which myosin II activity within the actin network (hereafter referred to as the ‘acto-myosin’ network) on NE remnants causes its contraction and CSV reduction after NEBD.

### The CSV reduction in prometaphase by acto-myosin network contraction ensures timely chromosome congression and anaphase onset

Following NEBD, spindle MTs interact with chromosomes while contraction of the NE-associated acto-myosin network reduces the CSV (Figure 1B), which would shorten the overall distance between chromosomes and spindle poles. Thus, acto-myosin network contraction might facilitate the interaction between chromosomes and spindle MTs, which would accelerate subsequent chromosome congression to the middle of the mitotic spindle. To test this, we measured the period between NEBD and completion of chromosome congression in cells expressing either LINC-DN or its control. The LINC-DN expression led to a considerable delay in chromosome congression and anaphase onset, compared with the control (Figure 4A), which was not associated with significant defects in spindle positioning or length (Figure S4A). In cells expressing LINC-DN, chromosomes often remained behind a spindle pole and outside of the pole-to-pole region after NEBD (Figure 4A; back-of-spindle chromosomes), which is consistent with the NE extension beyond spindle poles when contraction of the acto-myosin network was suppressed (Figure S3D). Back-of-spindle chromosomes still remaining at 20 min after NEBD in cells expressing LINC-DN led to a larger delay in congression (Figure S4B). Collectively, these results suggest that the CSV reduction due to contraction of the NE-associated acto-myosin network facilitates chromosome interaction with spindle MTs and diminishes back-of-spindle chromosomes, thus promoting timely chromosome congression and anaphase onset.

**Figure 4.**
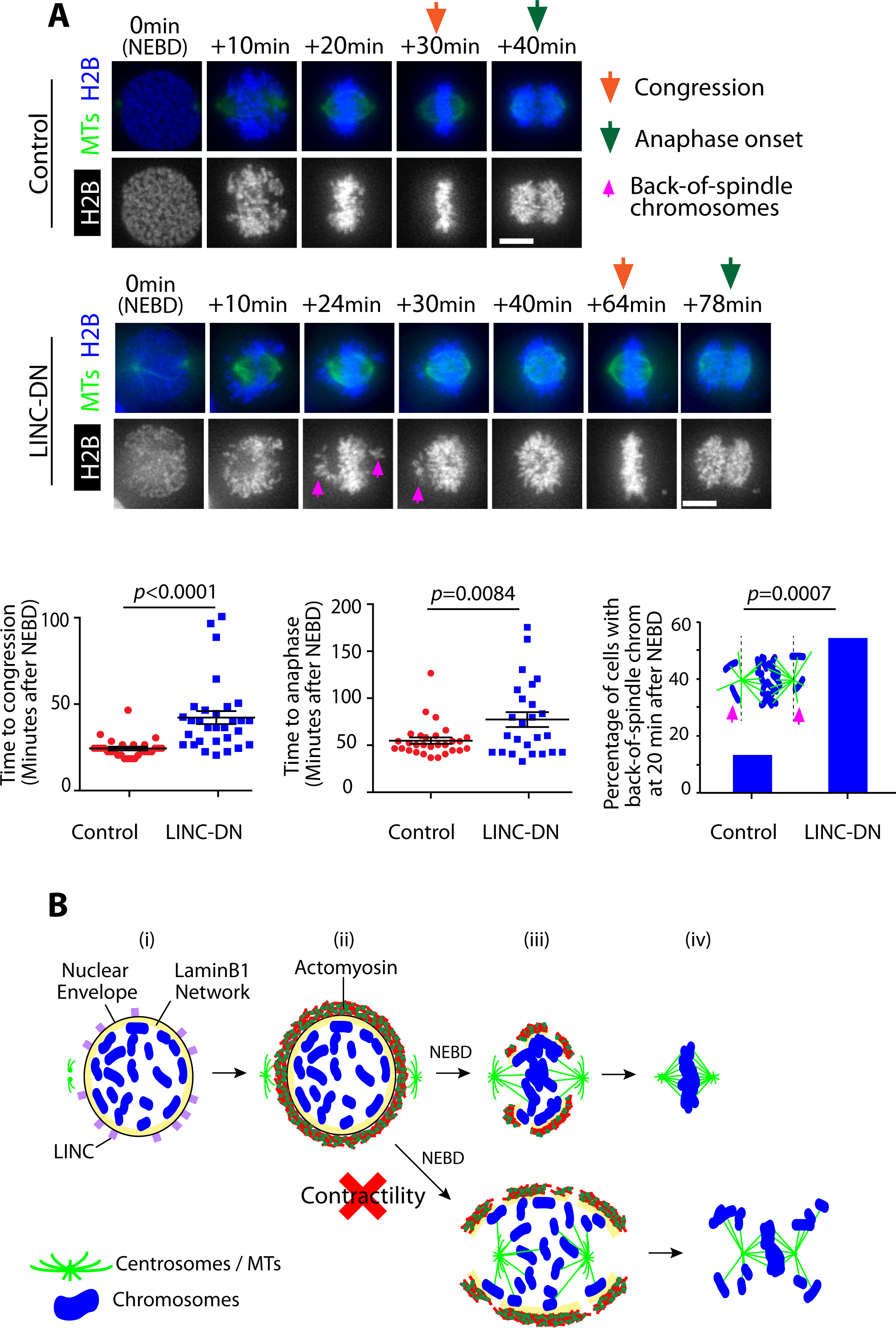
The CSV reduction in prometaphase by acto-myosin network contraction ensures timely chromosome congression and anaphase onset. A) Removal of actin network results in delays in congression and anaphase onset. Images (z-projections) show representative cells expressing histone 2B (H2B)-cerulean and GFP-tubulin. They progress from NEBD to anaphase, expressing either LINC-DN or a control construct. Orange arrows indicate time points at which congression was completed, while green arrows indicate time points of anaphase onset. Left-hand graph compares time (minutes) from NEBD to chromosome congression in cells expressing LINC-DN (n=30) versus a control construct (n=28). Middle graph compares time (minutes) from NEBD to the anaphase onset in cells expressing LINC-DN (n=25) versus a control construct (n=29). For these graphs, *p* value was obtained by *t*-test, error bars = s.e.m. Right-hand graph shows the percentage of cells with chromosomes at backside of the spindle at 20 min after NEBD in cells expressing LINC-DN (n=37) versus a control construct (n=31). *p* value was obtained by Fisher’s exact test. B) Summary diagram. Prior to NEBD, the LINC complex promotes the accumulation of acto-myosin to the cytoplasmic side of the nuclear envelope (i, ii). Upon NEBD, myosin II promotes contraction of the acto-myosin network on the NE remnant, thus reducing the CSV (iii). Constraining chromosomes from scattering results in efficient chromosome interaction with the spindle MTs as well as timely chromosome congression (iii, iv).

The above studies used human U2OS (near triploid) cells. We also investigated the acto-myosin network on the NE in other human cell lines. RPE cells (diploid) showed accumulation of actin on the NE in early mitosis (Figure S4C). By contrast, HeLa cells (highly aneuploid) did not show such NE-associated actin accumulation (Figure S4D). An interesting possibility is that a lack of the acto-myosin network is correlated with aneuploidy.

## Discussion

Chromosomes interact with spindle MTs during a narrow time window in early mitosis (i.e. prometaphase). To facilitate this process, the contraction of the acto-myosin network on NE remnants needs to occur immediately after NEBD. What triggers the contraction of the acto-myosin network at this specific timing in the cell cycle? We speculate that the reduction in nuclear membrane tension caused by NEBD may induce the contraction of the NE-associated acto-myosin network. Similar mechanisms have been reported (*14*); for example, during cell migration, reduced membrane tension within lamellipodia leads to local acto-myosin network contraction (*15*).

Recent studies have revealed important roles of the actin cytoskeleton in high-fidelity chromosome segregation. For example, in starfish oocytes, actin depolymerization beneath the NE facilitates the interaction of chromosomes with spindle MTs (*16, 17*). Moreover, in mouse oocytes, actin dynamics on the spindle promote robust kinetochore–MT interactions (*18*). In the current study, we report a further actin-dependent mechanism promoting the interaction between chromosomes and spindle MTs in mitotic human cells. In contrast to the two mechanisms mentioned above, in mitotic human cells an acto-myosin network resides on the cytoplasmic surface of the NE, and its contraction is driven by myosin II. Therefore, the actin cytoskeleton appears to facilitate efficient and robust chromosome interactions with spindle MTs through multiple mechanisms. It will be intriguing to explore how these mechanisms are conserved in evolution and used in different cell types to ensure correct chromosome segregation.

## Acknowledgements

We thank Tanaka lab members, A. Ciulli, A. Testa and M. Gierlinski for discussion; G. Ball for help with image analysis; L. Clayton for editing the manuscript; J. Januschke, T. Fukagawa, J. Swedlow, S. Megason, R. Adelstein, M. Davidson, and R.Y. Tsien for reagents; S. Swift, and P. Appleton for microscope maintenance. This work was supported by the Wellcome Trust (096535/Z/11/Z, 097945/Z/11/Z, 208401/Z/17/Z), Cancer Research UK (C28206/A114499) and Medical Research Council (MR/K015869/1). T.U.T. is a Wellcome Trust Principal Research Fellow. The authors declare no competing financial interests.

## Methods

### Cell Culture

To synchronize U2OS cells in early mitosis with a chemical genetic approach, their *CDK1* genes were replaced with *cdk1-as* construct (*4*), as previously done in DT40 cells (*19*). The U2OS *cdk1-as* cells, HeLa cells (ATCC CCL-2) and original U2OS cells (ATCC HTB-96) were cultured in DMEM (Gibco 41965-039) medium supplemented with 10% FBS (Gibco 10500) and 1% 10,000 U/ml Pen/strep (Gibco 15140-122). hTERT RPE (ATCC CRL-4000) were maintained in DMEM-F12 medium (Gibco 11330-057) supplemented with 10% FBS (Gibco 10500) and 1% 10,000 U/ml Pen/strep (Gibco 15140-122). To detach cells from culture flasks or dishes, they were treated with 0.05% Trypsin (Gibco 25300-054).

### Expression Constructs and transfection

Plasmid DNA constructs used in this study were: NLS-LacI-GFP (pT2699; (*20*)), Lifeact-mCherry (pT3118, a gift from Michael Davidson, Addgene #54491), GFP-Tubulin (pT2932, a gift from Jason Swedlow lab), LINC-DN (pT3138, mRFP1-SR-KASH; (*9*)) and LINC-DN Control (pT3139, mRFP1-KASH-ΔL; (*9*)); H2B-Cerulean (pT2698, a gift from Sean Megason lab), & MHC-GFP (pT3153, CMV-GFP-NMHC II-A, a gift from Robert Adelstein, AddGene #11347). For transfection of plasmids, reactions were prepared in 135µl OptiMem medium plus Glutamax (Gibco 51985-026) with 3µl DNA FuGene HD Transfection Reagent (Promega E2311) for every 1µg of plasmid DNA.

### Small molecule inhibitors

Small molecule inhibitors were used with the following concentrations: 1NM-PP1 1µM (Merck Millipore 529581), RO-3306 10µM (Merck Millipore 217699), nocodazole 3.3µM (Sigma M1404), paranitroblebbistatin (pnBB) 50µM (motorpharmacology), azidoblebbistatin (azBB) 5µM (motorpharmacology), jasplakinolide 1µM (Merck J4580) and MK-1775 500nM (Selleck Chemicals S1525). A Wee1 inhibitor MK-1775 was used to facilitate entry into mitosis by bypassing G2/M checkpoint (*21*) in our experiments with nocodazole (Figures 1C and 3B) and with LINC-DN (Figures 2C and 2D). Use of MK-1775 alone did not affect kinetics of CSV reduction.

### Microscopy

Live-cell images and immunofluorescence images were acquired using a Deltavision Elite widefield system (GE Healthcare) with 100× and 60× objective lenses (Olympus, NA 1.4) and cameras Cascade 1K EMCCD (Roper Scientific) and CoolsnapHQ2 (Photometrics). For live-cell imaging, the microscope chamber was maintained at 37°C with 5% carbon dioxide. For photoactivation of azBB, we used Zeiss 710 confocal system with a Coherent Chameleon multiphoton laser attachment and with 63× objective lens (NA 1.4). Super-resolution images were acquired using a Zeiss Airyscan 880 system with 63× objective lens (NA 1.4). Widefield Images were deconvolved using SoftWorx and images were analyzed using Fiji, OMERO, Imaris and Volocity software.

### Live-cell Imaging

Cells were synchronized and prepared for live-cell imaging as follows: cells were plated out at 50-60% confluency onto glass-bottomed imaging dishes (FluoroDish; World Precision Instruments, FD35-100). The following evening they were transfected with the relevant plasmids after replacement of the media with media lacking pen/strep. The transfection was allowed to continue overnight before the transfection reagents were washed out with further media without pen/strep. Cells were then arrested in the evening by addition of 1NM-PP1 (in the case of U2OS *cdk1-as* cells) or RO-3306 (in the case of HeLa or RPE cells), alongside SiR-DNA to stain chromatin where required. At this stage media was also replaced with Fluorobrite medium (Gibco A18967-01) supplemented with 10% FBS (Gibco 10500), 2mM L-Glutamine (Gibco 200mM, 25030-025), 25mM HEPES (1m Lonza BE17-737E) and 1mM Na-Pyruvate (100mM, Lonza BE13-115E). The following day cells were released from arrest by washing in 2 ml Fluorobrite medium (with supplements) 12 times, immediately followed by image acquisition. pnBB, azBB and MK-1775 were added after the final wash, prior to imaging. Nocodazole was added 1 hour before release, and added again after the final wash, prior to imaging. Images were acquired every 2 min except for Figure2A, Figure S2B and Figure S3D where images were acquired every minute, and Figure S1B where images were acquired every 5 min. For live-cell imaging of asynchronous U2OS cells, they were treated in the same way but were not arrested overnight before imaging. When a cell or a chromosome mass drifted during live-cell imaging, the frame of time sequence images in figures was shifted so that a chromosome mass was still positioned at the center of the frame in each image.

### Cell preparation for immunofluorescence

The night before fixation, media was replaced with Fluorobrite medium with supplements and the cells arrested with 1NM-PP1 if required. On the following day cells were released by washing in 2 mL Fluorobrite medium 12 times. Progression into mitosis was confirmed under a phase contrast light microscope. Then cells were fixed when they reached an appropriate stage, based on observation of chromosome condensation. To fix cells for immuno-staining, they were rinsed with pre-warmed PBS and incubated for 10 minutes with 37°C 4% methanol-free formaldehyde at 37°C. They were then rinsed 3× with PBS and permeabilized for 10 minutes in PBS with 0.5% Triton X-100. Following this they were blocked with 5% BSA in PBS for an hour at 37°C. While blocking, primary antibodies were prepared in 5% BSA in PBS to the relevant dilution. Following blocking, cells were incubated rocking with the primary antibody mixture for at least 60 mins at room temperature (or 4°C if incubated overnight). Following primary incubation, cells were washed rocking in PBS for 5 minutes, 3 times. Meanwhile, secondary antibodies plus phalloidin/Hoechst 33342 were prepared to the relevant dilution in 5% BSA in PBS. Cells were then incubated rocking with secondary antibodies for at least 60 mins at room temperature (or 4°C if incubated overnight). Following secondary incubation, cells were washed rocking in PBS for 5 minutes, 3 times.

### Immunofluorescence and cell staining

Primary antibodies were used for immuno-staining with the following dilution: anti-Tubulin 1:1000 (Cell Signalling Technology YL1/2), anti-LaminB1 1:1000 (Abcam ab16048) and anti-Actin 1:1000 (Thermo-Fisher PA1-183). Secondary antibodies were used with the following dilution: A488 goat anti-Rabbit 1:1000 (Thermo-Fisher A-11034), A488 donkey anti-Rat 1:1000 (Thermo-Fisher A-21208), and A594 goat anti-Rabbit 1:1000 (Thermo-Fisher A-11037). Dyes were used for cell staining with the following dilution: Phalloidin DyLight 650 1:1000 (Invitrogen 13454309), Hoechst 33342 1:2000 (Sigma-Aldrich 14533), SiR-DNA 100nm (tebu-bio SC007).

### Azidoblebbistatin photoactivation experiments

U2OS *cdk1-as* cells expressing NLS-LacI-GFP were prepared for live-cell imaging as described above. SiR-DNA was supplemented when 1NM-PP1 was added to arrest cells. Azidoblebbistatin (azBB) was added when cells were released from the arrest (by washing off 1NM-PP1). To identify cells in prophase, the start of chromosome condensation was monitored with SiR-DNA signal. A scan area was zoomed in so that it covered roughly half of the nucleus in prophase. A z-stack (3 sections, 2µm apart) of the region was then rapidly scanned four times with 633nm and 488nm lasers to observe chromosomes and NLS-LacI-GFP respectively, alongside 860nm multi-photon laser exposure to photoactivate the covalent binding of azBB to Myosin II. The region was scanned in three rounds in this way at 1-min interval. The scan area was then extended so that image of the entire nucleus could be acquired. The whole nucleus was then imaged for SiR-DNA and NLS-LacI-GFP in a z-stack of 20µm with 2µm z-interval, every 2 min. The timing of NEBD was judged based on the efflux of NLS-LacI-GFP from the nucleus into the cytoplasm. The nuclear region exposed to 860nm light could be discriminated based on partial photo-bleaching of SiR-DNA signals.

### Measurement of chromosome scattering volume

U2OS *cdk1-as* cells expressing NLS-LacI-GFP were imaged every 2 min in the presence of SiR-DNA. Z-stacks of 10-sections were acquired with 2µm interval. The CSV was measured as follows: The surface of chromosomes was identified using Imaris (3D surface object tool). Obviously non-chromosome objects were excluded in this process. On each z-section the surface contours of multiple chromosomes were connected into one single-stroke contour (Figure 1B, green lines imposed on images), using Imaris, in such a way that the connection doesn’t affect 2D convex hull. These contours were used to generate a continuous 3D surface contour, which was required to compute a 3D convex hull. Based on this continuous 3D surface contour, a 3D convex hull was generated using Imaris (convex hull generation tool). The volume of this 3D convex hull was defined as the CSV. To compare reduction of CSV between two conditions, we used two-way ANOVA in which data at all the time points shown in relevant graphs were included in analyses.

### Measurement of the volume and intensity of the actin network

U2OS *cdk1-as* cells expressing Lifeact-mCherry and GFP-tubulin were imaged every 2 min. Z-stacks of 10-sections were acquired at 2µm intervals. A 3D surface object was generated using Imaris by creating contours following the actin network at each z-section. The volume of the surface object was taken as the volume inside the actin network. Meanwhile, to measure the intensity of actin network, a 2 pixel wide line was drawn along the actin network on a single z-section using Fiji; the shape of the nucleus as seen by the exclusion of GFP-tubulin background was used as a guide when the actin signals were ambiguous. The mean gray value of the pixels within this line was taken as a measurement. As a background reading, a measurement was taken from a similar 2-pixel wide line drawn in the cytoplasmic region surrounding the nucleus. To generate a final reading the measurement from the cytoplasm was subtracted from the measurement from the actin network. The timing of NEBD was determined by the influx of GFP tubulin from the cytoplasm into the nucleus, judged by a sudden increase in GFP intensity within the nucleus.

### Measurement of chromosome mass volume

U2OS *cdk1-as* cells expressing NLS-LacI-GFP were imaged every 2 min in the presence of SiR-DNA. Z-stacks of 10 sections were acquired with 2µm interval. A 3D surface covering SiR-DNA signal was automatically generated with Imaris as follows; the mean value in the Cy5 channel was set as the threshold, and the Background Subtraction (local contrast) option was chosen. The volume of the surface object was defined as the chromosome mass volume (CMV). The efflux of NLS-LacI-GFP from the nucleus into the cytoplasm was taken to indicate the timing of NEBD.

### Timing of chromosome congression and anaphase onset

U2OS *cdk1-as* cells expressing GFP-tubulin and H2B-Cerulean were imaged every 2 min. Z-stacks of 10-sections were acquired with 2µm interval. The timing of NEBD was determined by the influx of GFP tubulin from the cytoplasm into the nucleus. Timing of complete congression was estimated as the time all chromosomes aligned on the spindle equator. Timing of anaphase onset was estimated as the time chromosomes began segregation to opposite poles.

## Supplemental Figures

**Figure S1.**
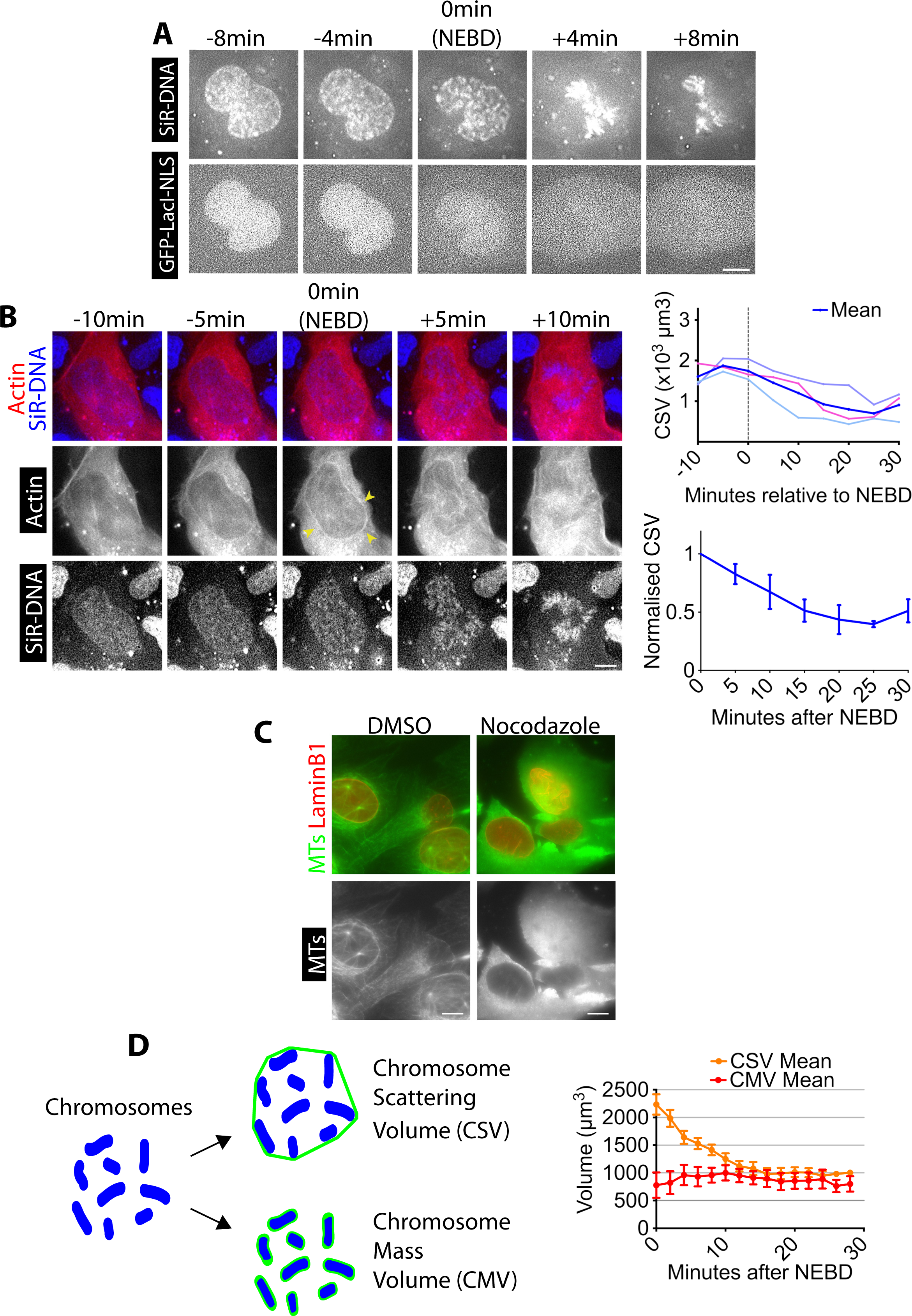
Supplemental data associated with Figure 1. A) Method for estimation of the NEBD timing. Images from the same time lapse as in Figure 1B. Top-row images show SiR-DNA-stained chromosomes. Bottom-row images show GFP-LacI-NLS (nuclear localization signal). The timing of NEBD was determined through observation of GFP-LacI-NLS spreading out of nucleus. Scale bars, 10µm. B) CSV decreases and actin accumulates on the NE around NEBD in asynchronous wild-type U2OS cells (without *cdk1-as*). Images show single z-sections of a U2OS cell with SiR-DNA-stained chromosomes (blue), expressing Lifeact-mCherry (red) and undergoing NEBD. Yellow arrowheads indicate location of the actin network. Time is shown, relative to NEBD. Timing of NEBD was determined as in Figure S1A. Scale bars, 10µm. C) Confirmation of the absence of MTs with nocodazole treatment. Immunostaining of fixed cells to test the presence or absence of microtubules (MTs) within cells. Cells were analyzed by live-cell imaging for Figure 1C, and then fixed and immunostained for MTs (and LaminB1 as a control) to confirm that MTs were depleted within cells, after nocodazole treatment (right) but not in control. Images are projections of 4 Z-sections at 1µm interval. Scale bars, 10µm. D) Chromosome mass volume did not considerably change after NEBD. Diagram (left) shows how CSV (chromosome scattering volume) and CMV (chromosome mass volume) were determined. Graph (right) shows the mean of CMV (n=3) and CSV (n=8). Error bars, s.e.m.

**Figure S2.**
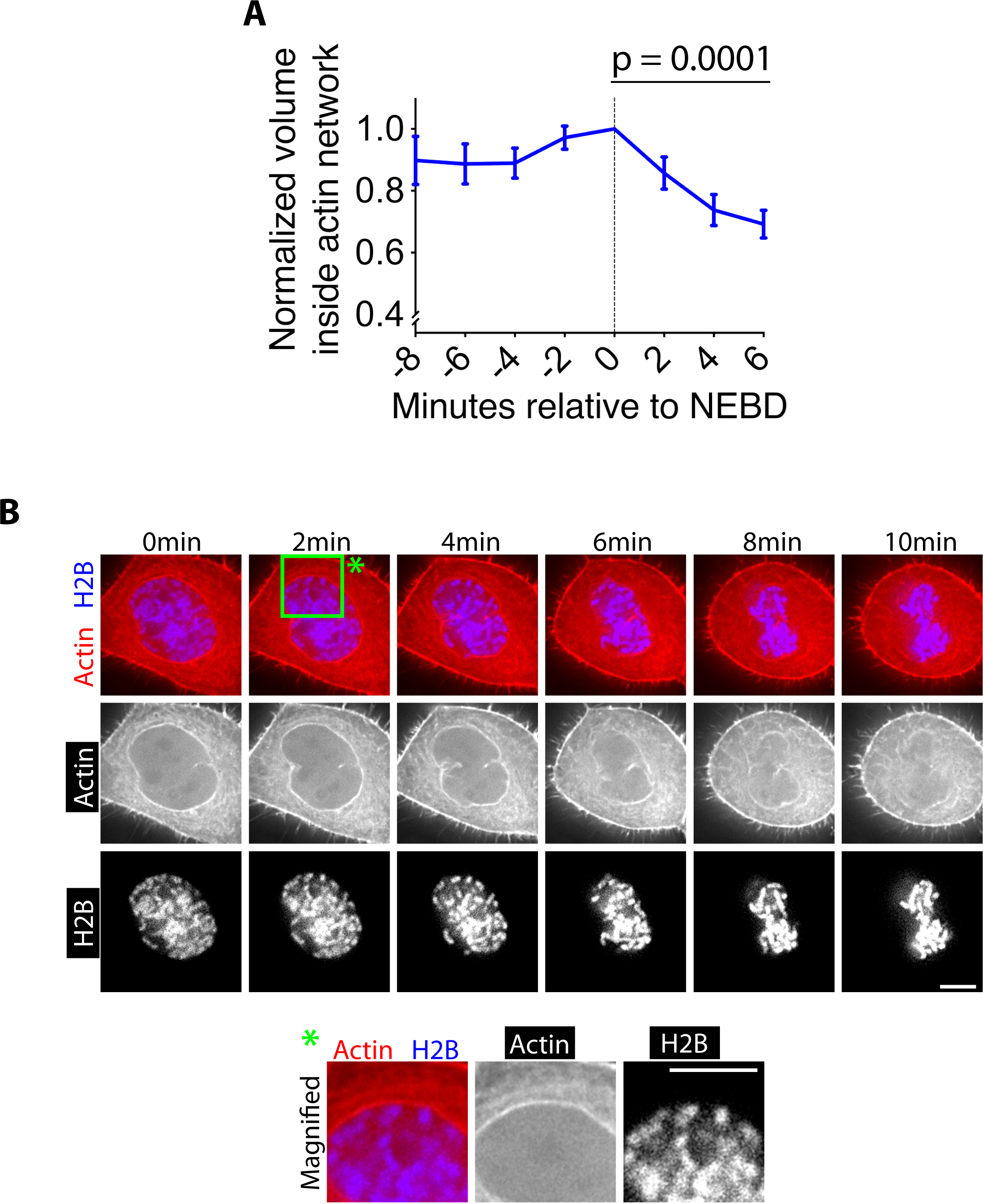
Supplemental data associated with Figure 2. A) The volume inside of the actin network is rapidly reduced after NEBD. Graph shows the mean of normalized volume inside the actin network (normalized to the volume at NEBD in each cell) after NEBD (n=7). Error bars, s.e.m. *p*-value was determined by one-way ANOVA for data during 0–6min. Note that it was difficult to quantify this volume after 6 min because actin signals were not continuous on the NE remnant region. B) During contraction of the actin network, outermost chromosomes are located right beneath the network. Images show single z-sections of a representative cell, expressing histone 2B (H2B)-cerulean and mCherry-Lifeact, in early mitosis. Green rectangle at 2 min indicates the region magnified in the images at bottom. Time 0 is set arbitrarily. Scale bars, 10µm.

**Figure S3.**
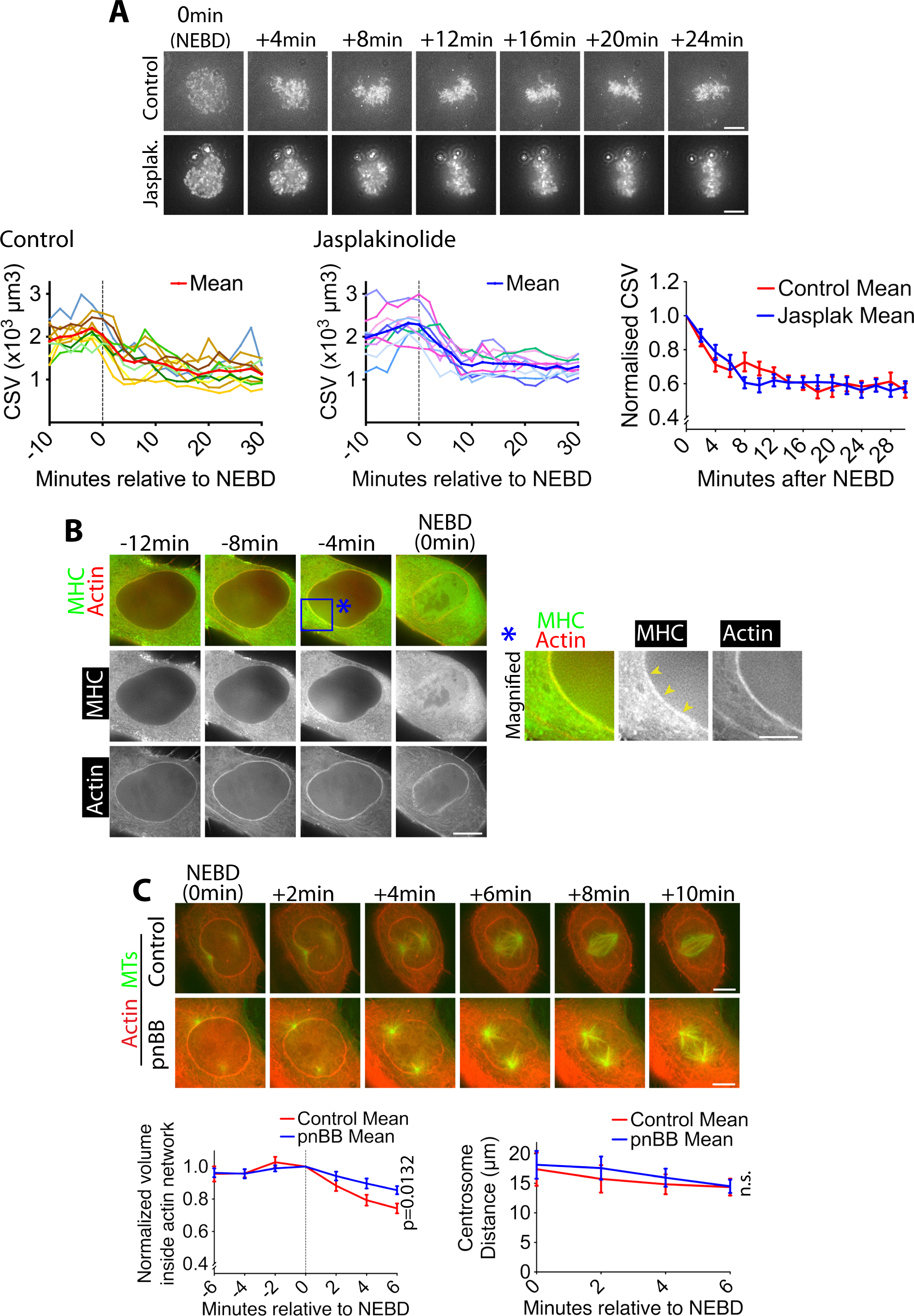
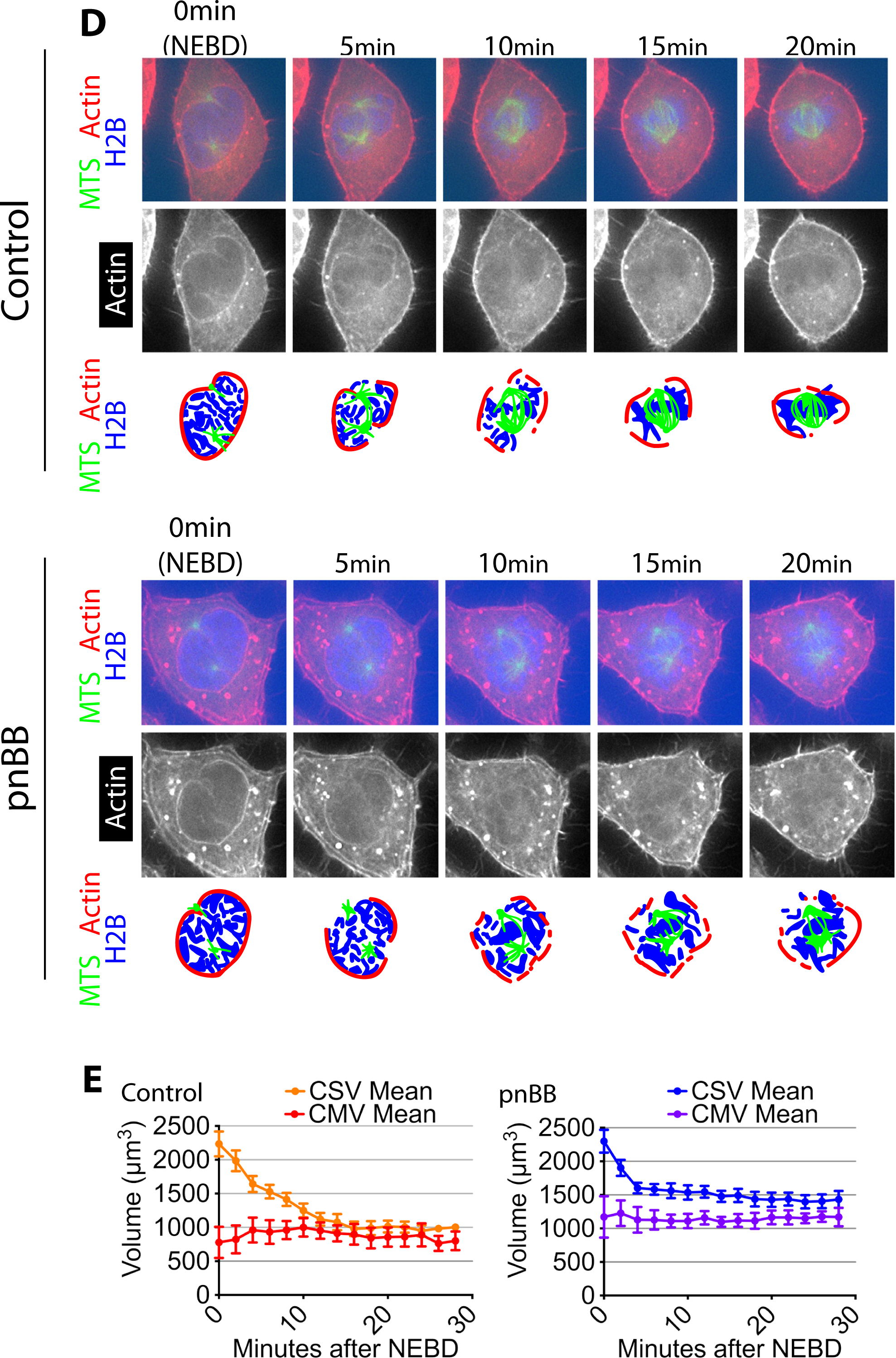
Supplemental data associated with Figure 3. A) Actin depolymerization inhibitor does not change CSV reduction after NEBD. Images (z-projections) show SiR-DNA-stained chromosomes in representative cells incubated in the presence of 1µM jasplakinolide (Jasplak., bottom) or DMSO (control, top). Time is relative to NEBD, which was determined as in Figure S1A. Scale bars, 10µm. Left-hand and center graphs show CSV measurements in individual cells in the presence and absence of jasplakinolide, respectively (n=10 each). The right-hand graph compares the means of normalized CSV (as in Figure 1B). Error bars. s.e.m. n.s. = not significant (control vs jasplakinolide-treated), determined by two-way ANOVA. B) Myosin II co-localizes with the actin network on the NE in prophase. Images on left (single Z-sections) show a representative cell expressing mCherry-Lifeact and GFP-NMHCII-A (GFP-tagged Myosin II-A heavy chain). MHC denotes myosin II heavy chain. Scale bars, 10µm. Images on right show magnification of the rectangle region at −4 min in the image at left. Yellow arrowheads indicate MHC signals co-localizing with the NE-associated actin network. Scale bars, 5µm. Time is shown relative to NEBD, whose timing was determined by the influx of GFP-NMHC-II A into the nucleus. C) Myosin II inhibition alleviates contraction of the actin network on the NE remnant after NEBD. Images (projections of two consecutive z-sections where spindle poles were in focus) show representative cells expressing GFP-tubulin and Lifeact-mCherry in the presence of 50µM paranitroblebbistatin (pnBB, bottom) or DMSO (control, top). Scale bars, 10µm. Timing of NEBD was determined by influx of GFP-tubulin into the nucleus. Left-hand graph shows the mean of normalized volume inside the actin network (normalized to the volume at NEBD in each cell) after NEBD in the presence (bottom, pnBB) and absence (top, control) of pnBB (n=8 each). *p* value (control vs pnBB) was obtained by two-way ANOVA. Error bars, s.e.m. Right-hand graph shows mean centrosome distance at the time points around NEBD in the presence (bottom, pnBB) and absence (top, control) of pnBB. n=8 in both conditions. *p* value (control vs pnBB) was obtained by two-way ANOVA. n.s., not significant. Error bars, s.e.m. D) Myosin II inhibition often leaves the actin network extending beyond the spindle poles soon after NEBD. Images (projections of two consecutive z-sections where spindle poles were in focus) show single z-sections of representative cells expressing GFP-tubulin, Lifeact-mCherry and H2B-cerulean in the presence of 50µM paranitroblebbistatin (pnBB, bottom) or DMSO (control, top). Scale bars, 10µm. Diagrams indicate position of actin network, chromosomes and MTs/ spindle poles. Timing of NEBD was determined by influx of GFP-tubulin (not incorporated to MTs) into the nucleus. In the example of a pnBB-treated cell shown here, the actin network extends beyond a spindle pole (on top) at 5 min after NEBD. E) Myosin II inhibition does not affect chromosome mass volume (CMV) after NEBD. Graphs show means of CSV and CMV in the presence (bottom, pnBB) and absence (top, control) of 50µM pnBB (CSV n=10, CMV n=3; error bars. s.e.m.). Graph at left-hand is identical to the graph in Figure S1C and shown here again for comparison.

**Figure S4.**
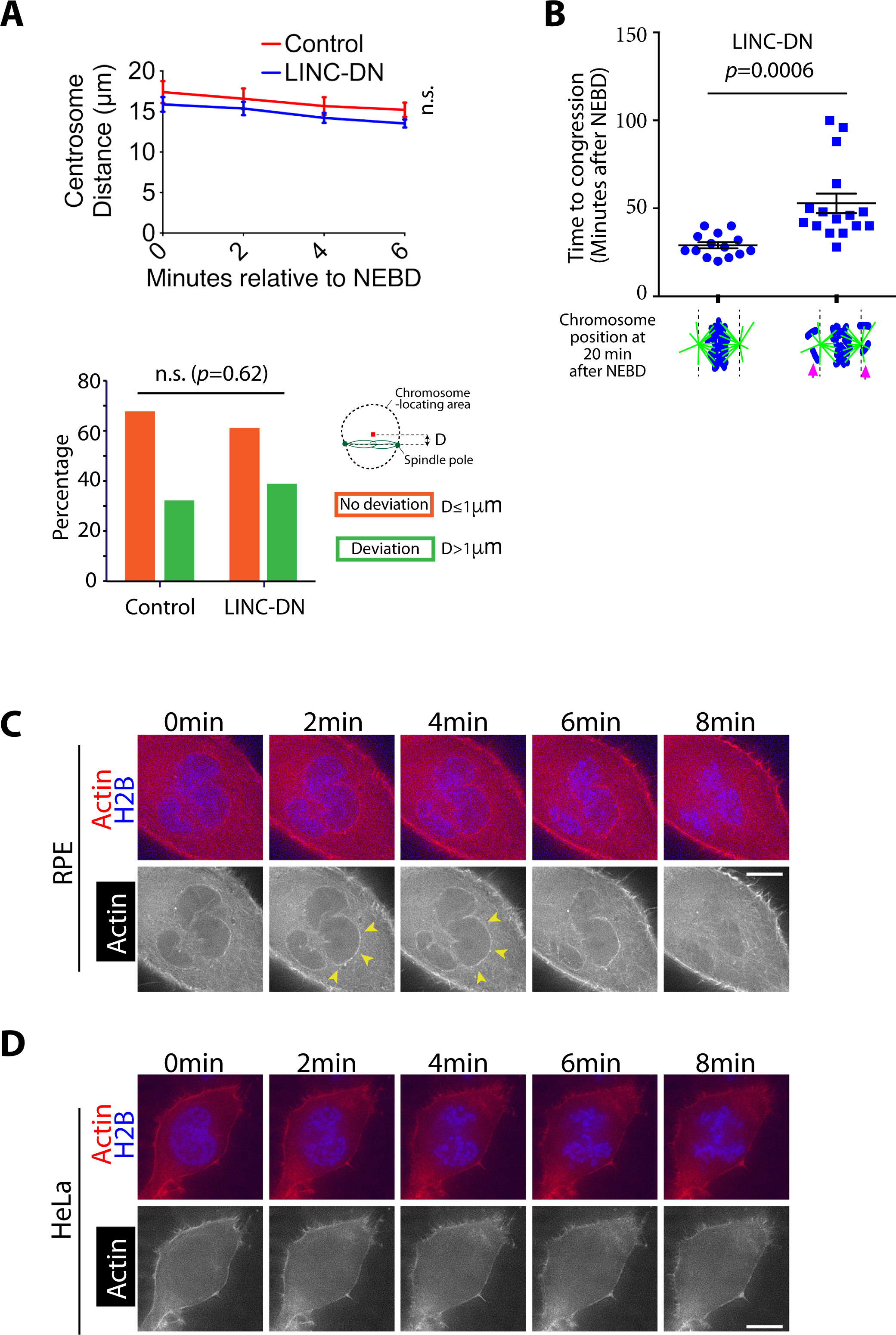
Supplemental data associated with Figure 4. A) LINC-DN expression did not change the distance between centrosomes and did not cause deviation of the spindle from the center of the nucleus. Images acquired for Figure 4A were analyzed further. Top graph shows the mean centrosome distance after NEBD in cells expressing a control construct (n=33) or LINC-DN (n=35). Error bars, s.e.m. *p* value (control vs LINC-DN) was obtained by 2-way ANOVA. n.s., not significant. Bottom graph shows the deviation of the spindle from the center of the chromosome-locating area, at 10 min after NEBD, in cells expressing a control construct (n=31) or LINC-DN (n=35). When the spindle axis is more (or less) than 1 µm from the center of the chromosome-locating area, it was scored that the spindle axis deviated (or not deviated) from the centre. n.s., not significant, determined by Fisher’s exact test. B) The presence of back-of-spindle chromosomes soon after NEBD leads to a delay in chromosome congression. Graph compares time (minutes) from NEBD to chromosome congression in cells expressing LINC-DN with (right, n=16) versus without (left, n=14) back-of-spindle chromosomes (magenta arrows) at 20 min after NEBD. Images acquired for Figure 4A were analyzed further. *p* value was obtained by *t*-test. C) RPE cells show accumulation of actin on the NE in early mitosis. Images (single Z-sections) show RPE cells expressing mCherry-Lifeact and H2B-cerulean, in early mitosis. Time 0 is set arbitrarily. Yellow arrowheads indicate location of actin network. Scale bars, 10µm. D) HeLa cells do not show apparent accumulation of actin on the NE in early mitosis. Images (single Z-sections) show HeLa cells expressing mCherry-Lifeact and H2B-cerulean, in early mitosis. Time 0 is set arbitrarily. Scale bars, 10µm.

